# A Syx-RhoA-Dia1 signaling axis regulates cell cycle progression, DNA damage, and therapy resistance in glioblastoma

**DOI:** 10.1101/2021.12.03.469255

**Authors:** Wan-Hsin Lin, Ryan W. Feathers, Lisa M. Cooper, Laura J. Lewis-Tuffin, Jann N. Sarkaria, Panos Z. Anastasiadis

## Abstract

Glioblastomas (GBM) are aggressive tumors that lack effective treatments. Here, we show that the Rho family guanine nucleotide exchange factor Syx promotes GBM cell growth both in vitro and in orthotopic GBM patient-derived xenografts. Growth defects upon Syx depletion are attributed to prolonged mitosis, increased DNA damage, G2/M cell cycle arrest, and cell apoptosis, mediated by altered mRNA and protein expression of various cell cycle regulators. These effects are phenocopied by depletion of the Rho downstream effector Dia1 and are due at least in part to increased cytoplasmic retention and reduced activity of the YAP/TAZ transcriptional coactivators. Further, targeting Syx signaling cooperates with radiation treatment and temozolomide (TMZ) to decrease viability in GBM cells irrespective of their inherent response to TMZ. Taken together, the data indicate that a Syx-RhoA-Dia1-YAP/TAZ signaling axis regulates cell cycle progression, DNA damage, and therapy resistance in GBM and argue for its targeting for cancer treatment.

**One Sentence Summary:** Syx promotes growth and therapy resistance in glioblastoma.

## Introduction

Glioblastomas (GBM) are malignant grade IV gliomas and are the most common primary brain tumors in adults. The hallmarks of this disease include aggressive growth and diffuse infiltration. Current standard of care for GBM involves maximal surgical resection followed by radiation therapy and chemotherapy with temozolomide (TMZ)^1^. Despite aggressive multimodal therapy, clinical outcomes for GBM patients remain dismal. In the United States, the median survival for GBM patients is about 15 months from initial diagnosis, and the 5-year survival rate is only about 5.5%^2^. Major contributors to this poor outcome are the inevitable recurrence of the tumor, which is underlined by its infiltrative nature and the difficulty of complete surgical resection, tumor heterogeneity, and resistance to treatment. TMZ is a DNA alkylating agent that causes methylation of guanine and adenine nucleotides. The accumulation of unrepaired DNA lesions promotes G2/M cell-cycle arrest and cell death, especially in tumors with low or no expression of O^6^-methylguanine-DNA-methytransferase (MGMT), a DNA repair protein^3^. Tumors lacking MGMT promoter methylation and high MGMT expression, account for more than 50% of GBM and show inferior responses to TMZ chemotherapy^4^. Regardless of MGMT status, recurrence is universal in GBM, pointing to an urgent need for new therapeutic strategies.

The Rho family of small GTPases is comprised of more than 20 members, including the most characterized RhoA, Rac1, and Cdc42. Rho GTPases regulate cytoskeletal remodeling, cell polarity, migration, gene expression, and cell cycle progression, coordinating diverse cellular functions. Rho guanine nucleotide exchange factors (GEFs) facilitate exchange of bound GDP for GTP, promoting Rho GTPase activation^5^. The active GTP-bound GTPases interact with downstream effectors to transduce signals to direct biological responses. Alterations of these GTPases and their regulators and effectors have been found in GBM and lead to aggressive disease^6^.

Synectin-binding RhoA exchange factor (Syx, also known as Tech, GEF720, Plekhg5) is a RhoA GEF that regulates junction integrity, cell migration, angiogenesis, and neural differentiation^7–11^. Syx is expressed in endothelial cells, where its localization to the apical cell-cell junctions is regulated by the scaffold protein Mupp1 (and its paralogue Patj), VEGF signaling, and 14-3-3 protein binding^7, 12^. Membrane-associated Angiomotin (Amot) forms a complex with Syx and Patj/Mupp1 to activate RhoA at the leading edge to direct endothelial cell migration^9, 11^. Moreover, Syx is widely expressed in both conventional and patient-derived xenograft (PDX) GBM lines, promoting cell chemotaxis^8^. The effects of Syx on junction integrity and cell migration are mediated by its ability to selectively induce the activation of mammalian Diaphanous formin (Dia1)^7, 8^.

RhoA and its effectors regulate gene expression and cell proliferation through multiple mechanisms, including the activation of the transcriptional cofactors Yes-associated protein 1 (YAP) and its paralog TAZ. Due to lack of a DNA-binding domain, YAP/TAZ associate with other transcription factors, such as TEADs, to direct downstream responses to promote organ growth and cell transformation^13, 14^. In GBM, YAP/TAZ expression and activity are thought to promote glioma aggressiveness^15, 16^. Activation of Hippo kinases (MST1/2, LATS1/2) in response to extracellular cues, such as cell confluency and cell adhesion, results in phosphorylation, nuclear exclusion, and proteasomal degradation of YAP/TAZ^13, 14^. Interestingly, expression of Amot and RhoA-mediated cytoskeleton remodeling can modulate YAP/TAZ activity via both Hippo kinase dependent and independent mechanisms^17, 18^. Moreover, activation of Dia1 promotes the function of YAP and TAZ^19, 20^.

Based on the roles of RhoA signaling in cell proliferation and transformation, we hypothesized that the previously established Syx-RhoA signaling might regulate GBM cell growth in addition to promoting directed cell migration. Using conventional and xenograft GBM lines, we show here that downregulation of Syx-RhoA-Dia1 signaling in GBM cells results in cell cycle arrest, mitotic failure, DNA damage and increased apoptosis *in vitro*, as well as increased overall survival in orthotopic GBM xenograft bearing mice. These effects are mediated, at least in part, by impaired activity of YAP/TAZ, leading to deregulated expression of key cell cycle regulators. Importantly, targeting this pathway augments cell responses to TMZ and radiation, pointing to its potential utility in GBM therapy.

## Results

### Depletion of Syx decreases GBM cell growth

Increased proliferation and tumor cell dissemination are major factors in the poor prognosis of GBM patients. We reported previously that Syx is expressed in GBM cells and is required for directed cell migration^8^. To examine whether Syx also affects GBM cell growth, we utilized several conventional (U251, LN229) and PDX lines (GBM10, GBM12, GBM14). GBM10, GBM12, and GBM14 lines represent distinct transcriptional subtypes of GBM (GBM10 and GBM12 are mesenchymal and GBM14 is classical subtype) and respond differently to TMZ due to their methylation status of the MGMT promoter (GBM12 cells are MGMT methylated and sensitive to TMZ, whereas GBM10 and GBM14 cells are MGMT unmethylated and TMZ-resistant^21^). Knockdown of *Syx (Plekhg5)* by each of two non-overlapping shRNAs resulted in a significant decrease in cell growth in all tested GBM lines (Fig. 1a-c and Fig. S1). Next, we assessed the effect of *Syx* knockdown on GBM growth and overall survival *in vivo* using the orthotopic xenograft model. GBM12 cells were infected with lentiviruses expressing different shRNAs, as well as with a luciferase-expressing virus. Infected cells were then implanted intracranially. Tumor growth was monitored using bioluminescence imaging. *Syx* knockdown resulted in decreased tumor growth along with increased overall survival compared to the NT-sh control (Fig. 1d, e). Expression of *Syx* RNA transcripts was assessed using RNAscope analysis to confirm the knockdown *in vivo* (Fig. 1f). Overall, the data argue that depletion of Syx inhibits GBM cell growth *in vitro* and *in vivo*.

**Fig. 1.**
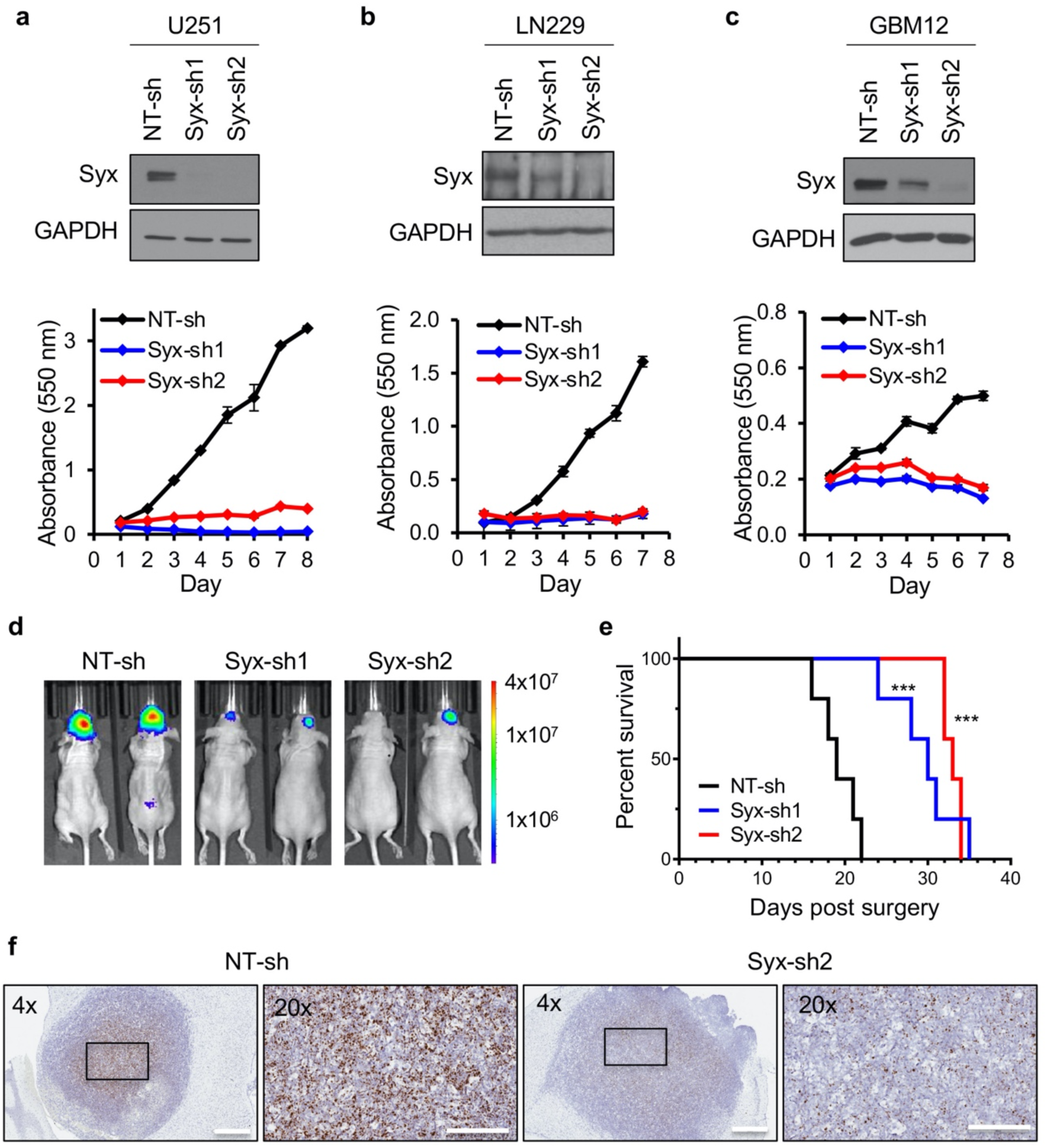
Depletion of Syx decreases GBM cell growth. **a-c** Immunoblot analysis of Syx and GAPDH in lysates from GBM conventional [U251 (**a**), LN229 (**b**)] and PDX [GBM12 (**c**)] cell lines transduced with indicated shRNAs (top). Cell viability over indicated time for each cell population was measured by the MTT assay (bottom). Shown are representative graphs with three technical replicates of at least three biological repeats. Graphs represent the means ± SD. **d** Representative images of brain bioluminescence on day 19 post-transplantation from intracranial xenografts derived from GBM12 cells expressing indicated shRNAs in immunocompromised mice. **e** Kaplan-Meier survival curves of mice orthotopically transplanted with GBM12 cells transduced with indicated shRNAs. *n* = 5 mice per group. Log-rank test, (\*\*\**P* < 0.001 for either Syx-sh1 or Syx-sh2 compared to NT-sh). **f** Expression of human Syx transcripts detected by RNAscope in situ hybridization in GBM12-derived xenografts expressing indicated shRNAs. Scale bar, 600 μm (4x), 200 μm (20x).

### Syx is required for GBM cell cycle progression and mitosis

To investigate how Syx depletion affects cell growth, we examined its effects on cell cycle progression and cell apoptosis. Cell cycle analysis using flow cytometry and propidium iodide was performed in U251 cells expressing *Syx*-shRNAs or NT-shRNA control. Depletion of Syx resulted in cell cycle arrest at the G2/M phase compared to control (Fig. 2a). To test whether this reflected a blockage of G2-to-M transition leading to reduced mitotic entry, we examined the level of phosphorylated histone H3 at Ser10 (pHH3), a well-accepted mitotic marker^22^. Depletion of Syx resulted in reduced pHH3 levels in different GBM lines, suggesting a defect in mitotic entry (Fig. 2b). Moreover, the expression of two mitotic regulators, Cdc20 and Survivin, was also downregulated upon *Syx* knockdown (Fig. 2b and Fig. S2). Cdc20 is a key cofactor of the anaphase-promoting complex (APC/C) ubiquitin ligase, which controls separation of sister chromatids in the metaphase-to-anaphase transition, as well as mitotic exit^23^. During mitosis, Survivin, a member of the chromosome passenger complex (CPC), governs proper chromosome positioning and segregation^24^. To directly examine cell division, we performed time-lapse imaging of cells expressing fluorescently labelled histone H2B to better visualize chromosomes (Fig. 2c-e and Movies S1 to S3). During the 24-hour imaging period, approximately 95% of NT-shRNA cells were able to enter and complete cell division (Fig. 2c). Consistent with the reduced pHH3 levels (Fig. 2b), only 6% of *Syx*-sh1 and 22% of *Syx*-sh2 cells were able to undergo successful cell mitosis (Fig. 2c). Furthermore, NT-sh cells completed mitosis in approximately 1.7 hours, whereas the *Syx* knockdown cells that were able to undergo mitosis required at least 5 hours (Fig. 2d). Combined, the data suggest that Syx depletion results in two phenotypes, a predominant defect in mitotic entry, as well as prolonged mitosis.

**Fig. 2.**
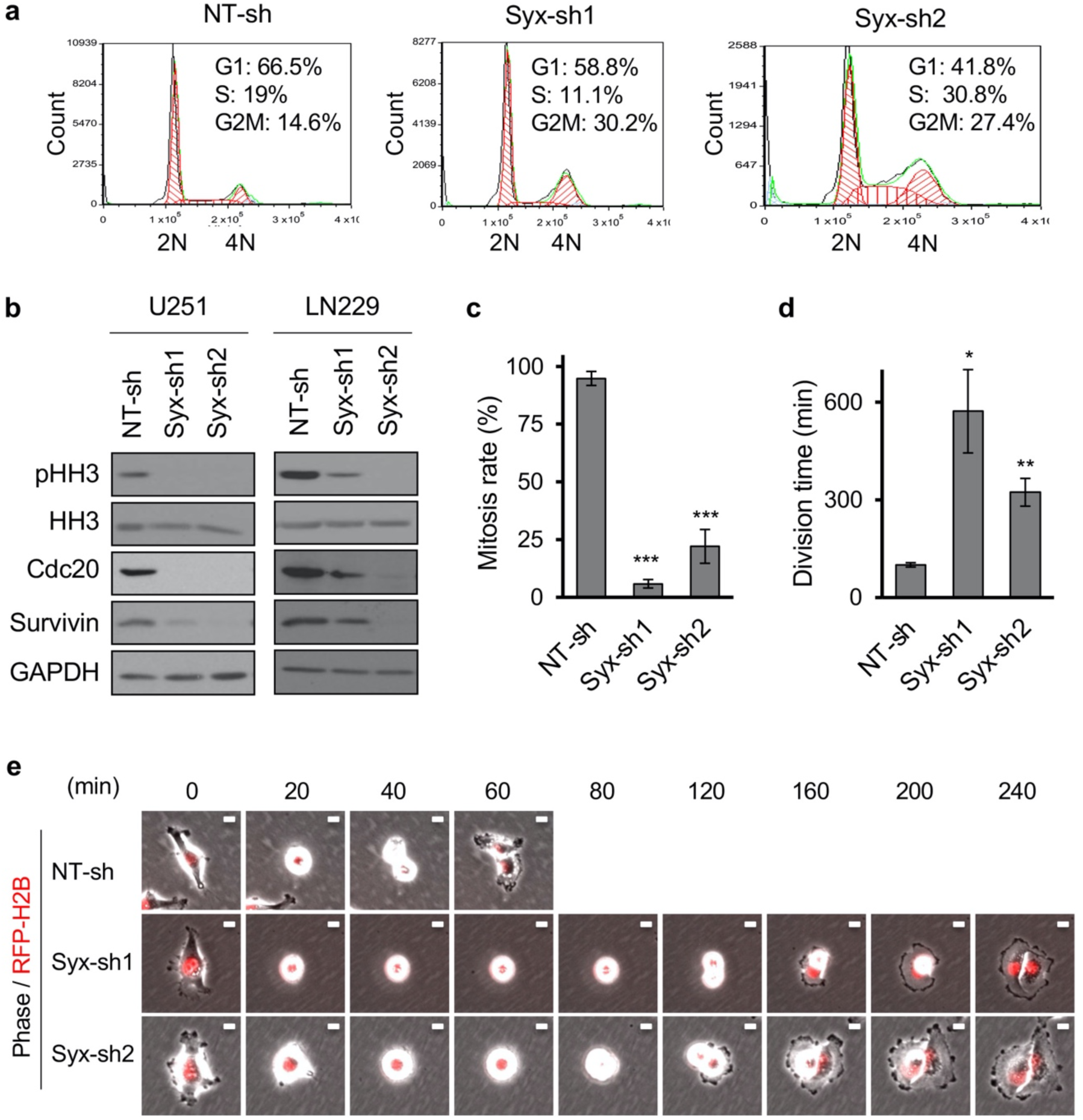
Syx is required for GBM cell cycle progression and mitosis. **a** DNA fluorescence histograms and percentages of cells at different cell cycle phases, as determined by propidium iodide-based DNA cell cycle analysis. DNA content (2N, 4N) is indicated. **b** Immunoblot analysis of mitotic markers (phosphorylated histone H3 at Ser10, pHH3), total histone 3 (HH3) and mitotic regulators (Cdc20, Survivin) in lysates of GBM cells (U251, LN229) expressing indicated shRNAs. **c**-**e** Cell division was visualized and analyzed by 24 hr time-lapse imaging of U251 cells expressing indicated shRNAs and RFP-H2B. **c** Percentage of cells with successful cell division in each group (*n* > 100 cells per group). **d** The duration of mitosis (division time) of cells that successfully underwent mitosis. Graphs (**c-d**) represent the mean ± SEM of 3 biological repeats. Student’s *t* test, \**P* < 0.05, \*\**P* < 0.01, \*\*\**P* < 0.001. **e** Representative images of cells undergoing mitosis, acquired by time-lapse microscopy (time in minutes indicated above). Scale bar, 10 μm.

### Syx depletion increases DNA damage

Prolongation of mitosis has been linked to double-stranded DNA damage and activation of the DNA damage response (DDR)^25, 26^, in part through the activation of executioner caspases (caspase 3/7). Consistent with this, knockdown of *Syx* resulted in upregulation of cleaved caspase 3 and poly(ADP-ribose) polymerase (PARP), accompanied by downregulation of un-cleaved PARP (Fig. 3a). Additionally, incubation of U251 cells with a Cas3-7 biosensor, which becomes fluorescent upon activation of executioner caspases, indicated that more *Syx* knockdown cells exhibit emission of fluorescent signal compared to control NT-sh cells (Fig. 3b).

**Fig. 3.**
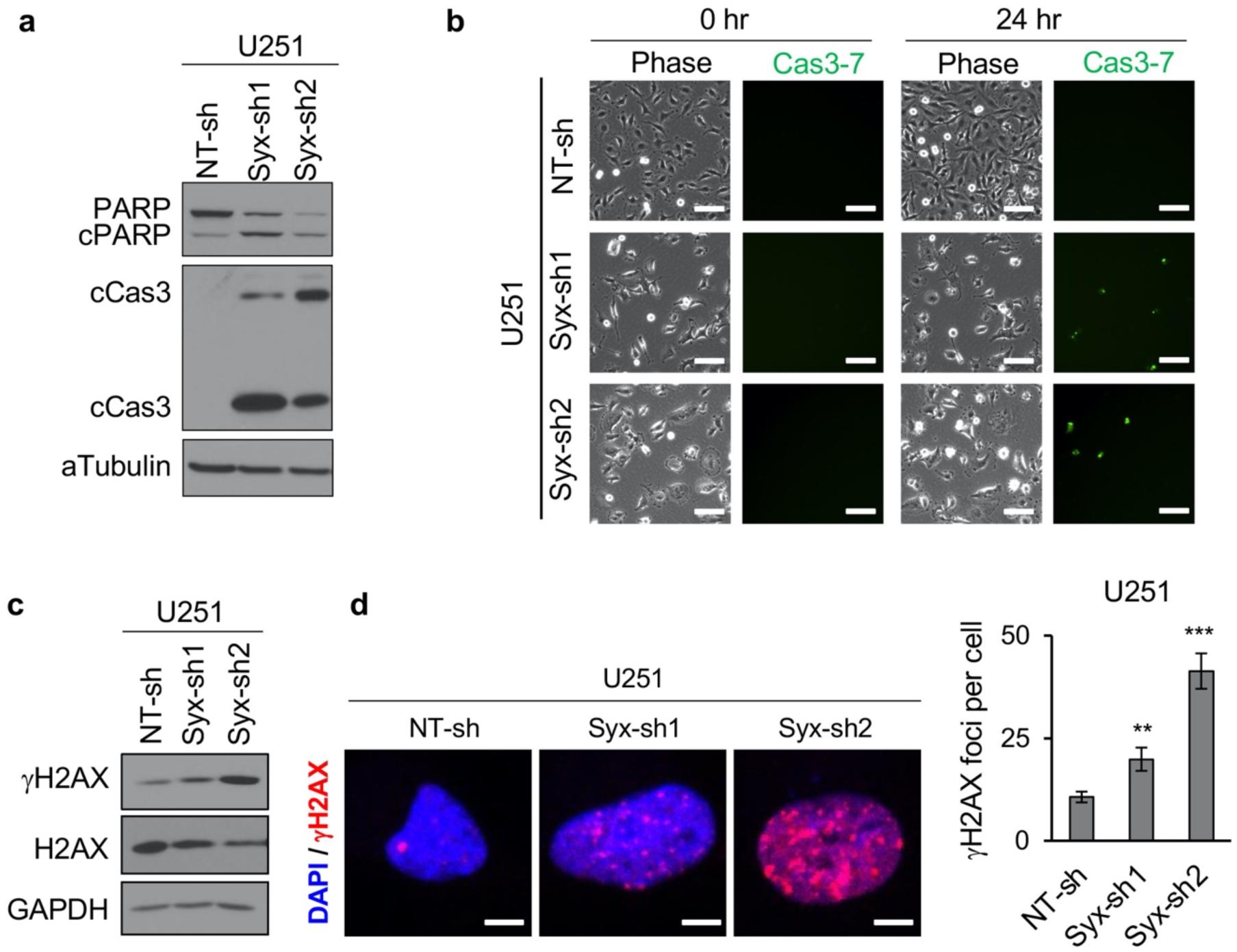
Syx depletion increases DNA damage. **a** Immunoblot analysis of apoptotic markers [cleaved caspase 3 (cCas3) and cleaved PARP (cPARP)] and *α*-tubulin in lysates of U251 cells transduced with Syx shRNAs. **b** Phase contrast (Phase) and corresponding fluorescence images of activated effector caspases (caspases-3/7) using a Cas3-7 probe. Shown are images acquired at time 0 and 24 h after addition of the Cas-3/7 probe. Scale bar, 100 μm. **c** Immunoblot analysis of phosphorylated H2AX at Ser-139 (*γ*H2AX), total H2AX, and GAPDH in lysates of U251 cells expressing indicated shRNAs. **d** Immunofluorescence staining of nucleus (DAPI, blue) and *γ*H2AX foci (red) in U251 cells expressing indicated shRNAs. Representative images are shown (left). Scale bar, 5 μm. Bar graph (right) depicts the average ± SEM number of *γ*H2AX foci per cell in U251 cells transduced with indicated shRNAs (*n* > 50 cells per group). Student’s *t* test, \*\**P* < 0.01, \*\*\**P* < 0.001.

To test if DNA damage is induced upon Syx depletion, we assessed the level of Ser139-phosphorylated Histone H2AX (known as *γ*H2AX), a sensitive marker of double strand breaks (DSBs)^27^. Depletion of Syx increased both the phosphorylation of H2AX and the overall number of *γ*H2AX nuclear foci in U251 cells (Fig. 3c, d). Importantly, similar results were also obtained in GBM10 and GBM12 PDX lines (Fig. S3a-d).

Previously, we and others reported a crucial role of Syx in the regulation of endothelial cell junction stability, vascular permeability, and endothelial cell migration^7, 9, 11^. To account for non-tumor specific effects of inhibiting Syx, we assessed whether Syx silencing also induces DNA damage in primary human brain microvascular endothelial cells (HBMECs). In contrast to GBM cells, Syx depletion did not increase the number of *γ*H2AX foci in HBMECs (Fig. S3e, f).

### Syx regulates the expression of cyclins and cyclin-dependent kinase inhibitors

The transition from one phase of the cell cycle to another is regulated by the coordinated action of cyclins, cyclin-dependent kinases (CDKs), and CDK inhibitors (CKIs)^28^. Knockdown of *Syx* in either U251 or LN229 GBM cells resulted in upregulation of the CKIs p21^Cip1^ and p27^Kip1^, and downregulation of cyclin E2, cyclin A2, and cyclin B1 (Fig. 4). Similar effects were also observed in GBM PDX lines (Fig. S4a). Using the CDK1 inhibitor RO-3306 to synchronize cells in the G2 phase^28^, the level of pHH3 in NT-shRNA cells increased as cells entered mitosis upon RO-3306 release, while the level of cyclin B1 progressively decreased as cells exited mitosis (Fig. S4b). Of note, despite being arrested in G2/M (Fig. 2a), *Syx* knockdown cells exhibited very low expression of cyclin B1 both before and after release from RO-3306 treatment (Fig. S4c). This result suggests that deregulation of cyclin B1 (and possibly other cell cycle regulators) upon Syx deficiency is not a consequence but likely a cause of cell cycle arrest.

**Fig. 4.**
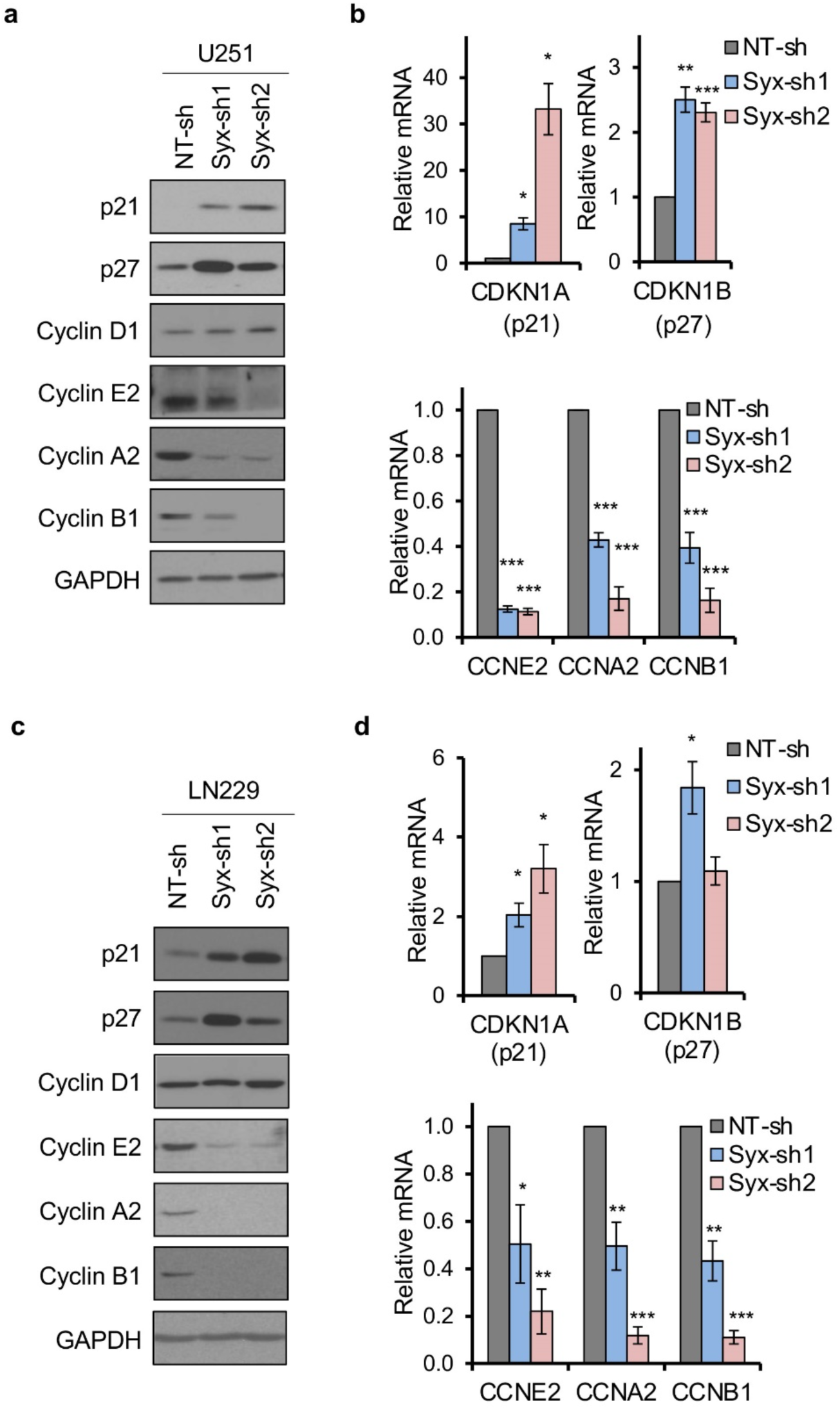
Syx regulates the expression of cyclins and cyclin-dependent kinase inhibitors. **a-d** Immunoblotting (**a**, **c**) and RT-qPCR (**b**, **d**) analyses of the expression of p21 (CDKN1A), p27 (CDKN1B), Cyclin D1, Cyclin E2 (CCNE2), Cyclin A2 (CCNA2), and Cyclin B1 (CCNB1) in U251 (**a**, **b**) and LN229 (**c**, **d**) cells expressing indicated shRNAs, grown at sub-confluency. Bar graphs (**b, d**) represent mean ± SEM of 3-4 biological replicates of relative mRNA expression of indicated genes normalized by GAPDH or *β*-actin. Student’s *t* test, \**P* < 0.05, \*\**P* < 0.01, \*\*\**P* < 0.001.

To identify the underlying mechanism of cyclin and CKI deregulation in Syx knockdown cells, we performed quantitative RT-PCR to measure the RNA levels of these molecules. The data indicated that deregulation of cyclins and CKIs in these cells occurs at the mRNA level (Fig. 4b, d). Interestingly, Cdc20 and Survivin transcripts were also downregulated upon *Syx* silencing (Fig. S2a, b). The results argue that Syx affects transcription to regulate cell cycle progression.

### Dia1 and YAP/TAZ signaling are downstream effectors of Syx

To gain insight into the mechanism of Syx action, we investigated the involvement of potential downstream effectors of the Syx-RhoA signaling pathway. First, we focused on the role of the formin Dia1, a known downstream effector of Syx in the context of endothelial junction formation and GBM cell migration^7, 8^. Similar to *Syx* knockdown, downregulation of *Dia1(Diaph1)* by each of two non-overlapping shRNAs^7, 8^, resulted in decreased cell growth in multiple GBM lines (Fig. 5a-c and Fig. S5a). Moreover, expression of cyclins, CKIs, and mitosis regulators was deregulated at both mRNA and protein levels (Fig. 5d, e). Downregulation of pHH3 levels suggested a defect in mitotic entry (Fig. 5d). Additionally, knockdown of *Dia1* increased DSBs as indicated by *γ*H2AX immunofluorescence staining (Fig. 5f). The ability of Dia1 to phenocopy Syx effects on the expression of cell cycle regulators, overall cell growth, and DNA damage, argues that Dia1 is a key downstream effector of Syx growth-related signaling.

**Fig. 5.**
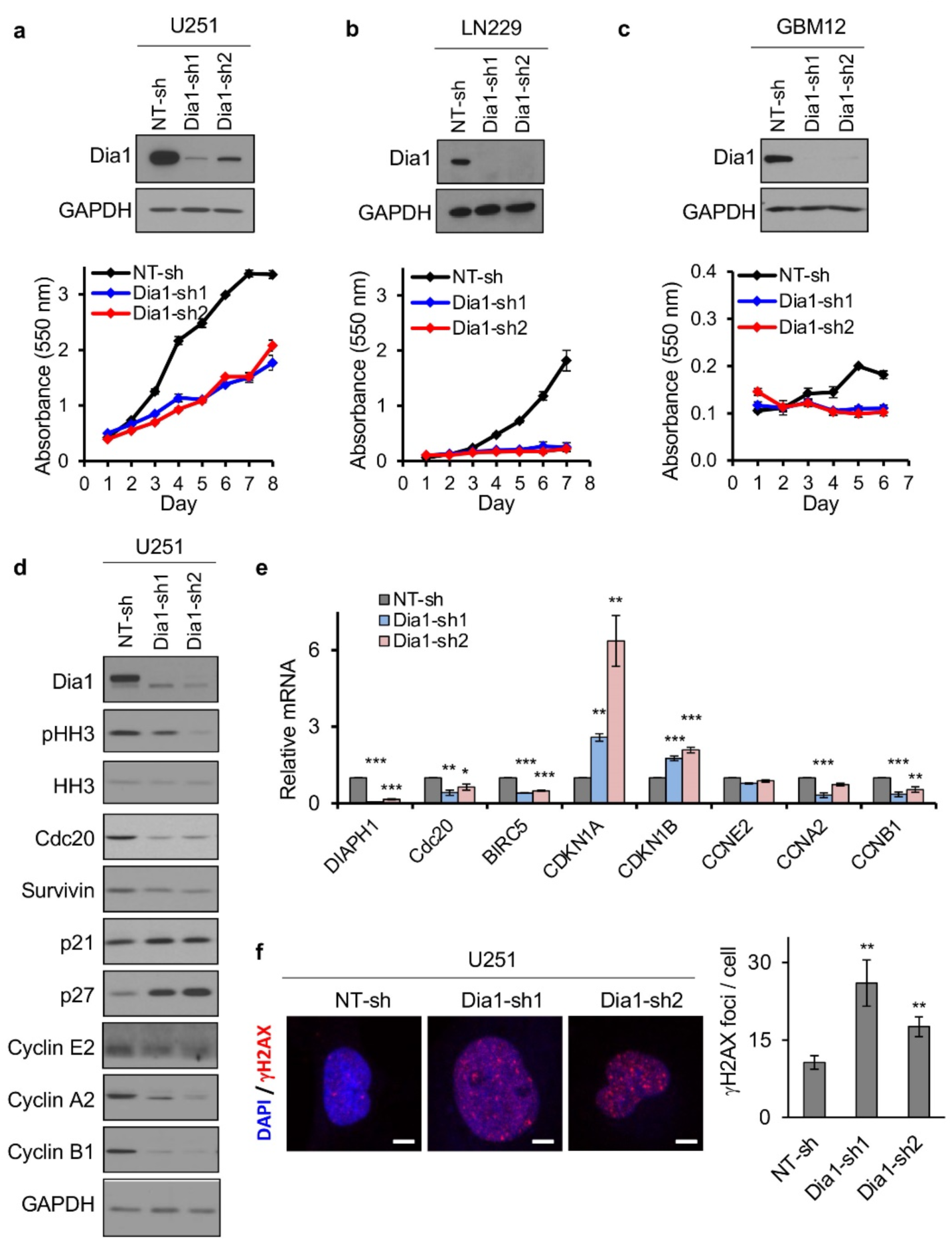
Dia1 is a downstream effector of Syx for the regulation of cell growth and gene expression. **a**-**c** Immunoblot analysis of Dia1 and GAPDH in lysates of U251 (**a**), LN229 (**b**), or GBM12 (**c**) cells transduced with *Dia1* shRNAs (top). Cell viability over indicated time for each cell population was measured by the MTT assay (bottom). Shown are representative graphs (means ± SD) of 3 biological repeats with 3 technical replicates each. **d, e** Immunoblot (**d**) and RT-qPCR (**e**) analyses of phosphorylated histone 3 at Ser-10 (pHH3), total histone 3 (HH3), Cdc20, Survivin (BIRC5), p21 (CDKN1A), p27 (CDKN1B), Cyclin E2 (CCNE2), Cyclin A2 (CCNA2), Cyclin B1 (CCNB1) in U251 cells expressing indicated shRNAs. Graph (**e**) represents the mean ± SEM of more than three biological replicates of relative mRNA expression of indicated genes normalized by GAPDH. **f** Representative images of immunofluorescence staining (left) shows *γ*H2AX (red) foci in the nucleus (DAPI, blue) of U251 cells expressing indicated shRNAs. Scale bar, 5 μm. Bar graph (right) depicts the average ± SEM number of *γ*H2AX foci per cell in U251 cells transduced with indicated shRNAs (*n* > 60 cells per group). Student’s *t* test, \**P* < 0.05, \*\**P* < 0.01, \*\*\**P* < 0.001.

Next, we focused on signaling pathways downstream of Syx-RhoA-Dia1 that could influence the expression of cell cycle regulators. One pathway by which RhoA drives transcription involves serum response factor (SRF)^29^ and its co-activator myocardin-related transcription factor (MRTF), which couple changes in actin dynamics and cell shape to gene expression and cell growth^30^. Another pathway by which both RhoA and Dia1 affect transcription is through the regulation of YAP/TAZ activity^19, 20^. Interestingly, YAP/TAZ have been previously shown to regulate cell cycle progression^31, 32^, and the expression of cyclins, CKIs, Survivin, and Cdc20 in other cell types, either directly or indirectly^32–34^. Consistent with an involvement of YAP/TAZ in the Syx-RhoA-Dia1 signaling axis, silencing of either *Dia1* or *Syx* resulted in increased cytoplasmic localization of YAP/TAZ and concomitant decrease in nuclear localization relative to the NT-shRNA control (Fig. 6a, b and Fig. S5b, c). Moreover, expression of connective tissue growth factor (CTGF), a well-established direct target gene of YAP/TAZ^14^, was also downregulated by *Syx* silencing (Fig. 6c). Additionally, knockdown of *Syx* resulted in downregulation of YAP/TAZ activity by at least 2-fold (Fig. 6d), as measured by a YAP/TAZ-responsive luciferase reporter (GTIIC-Luc)^35^. In contrast, ectopic expression of Syx resulted in increased transcriptional activity of YAP/TAZ (Fig. 6e).

**Fig. 6.**
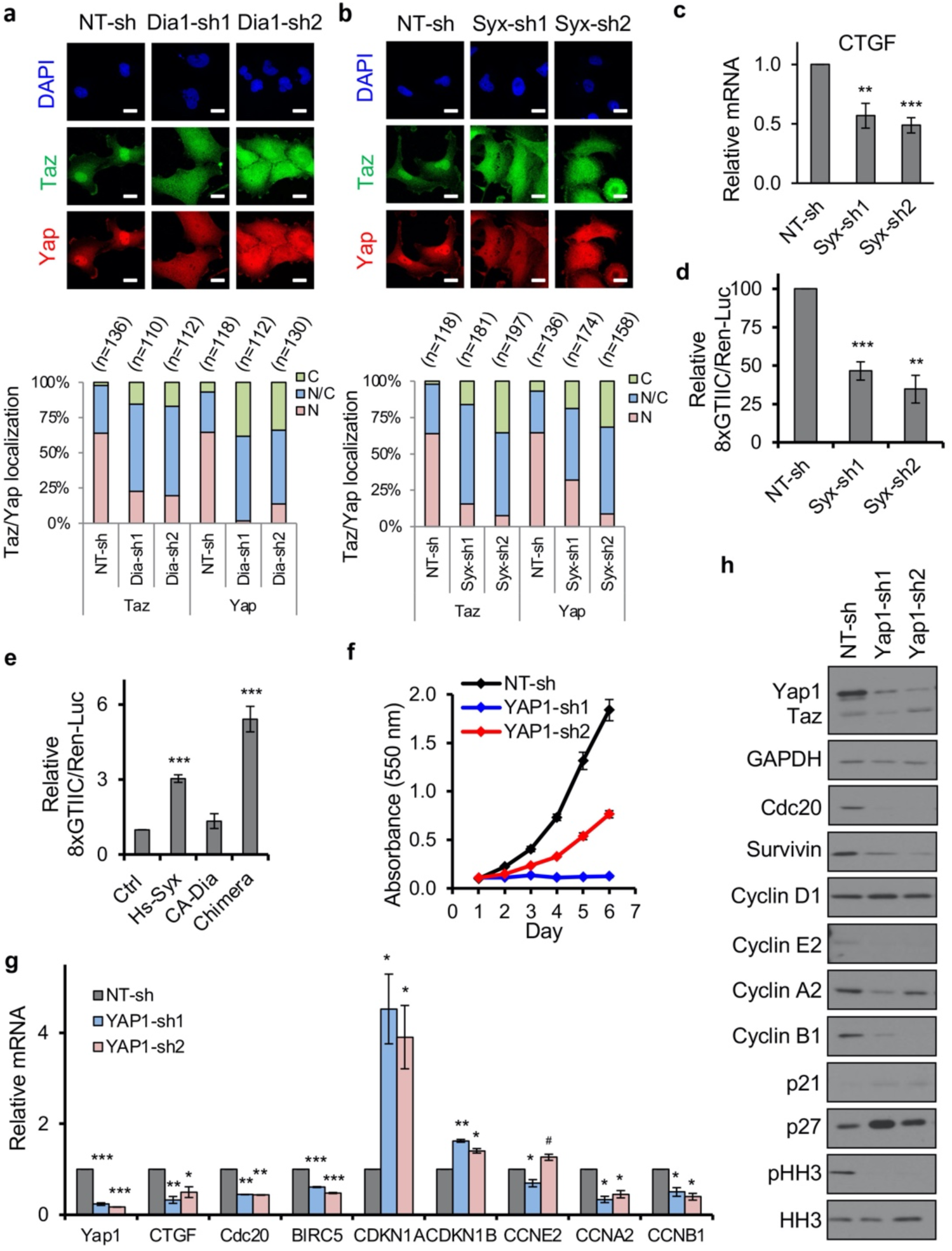
Dia1 and YAP/TAZ are downstream effectors of Syx. **a, b** Immunofluorescence images (top) of subcellular localization of YAP (red) and TAZ (green) in U251 cells expressing *Dia1* (**a**) or *Syx* (**b**) shRNAs. Scale bar, 15 μm. Staggered graphs (bottom) depict percentage of cells with YAP and TAZ in the cytosol (C), nucleus (N), or both (N/C). **c** RT-qPCR analysis of relative mRNA levels of CTGF in U251 cells expressing indicated shRNAs. Graph represents the mean ± SEM of 5 biological replicates. **d, e** Graphs show relative luciferase activity of a YAP/TAZ responsive reporter (8xGTIIC) in U251 cells expressing indicated shRNAs (**d**) or expressing indicated constructs (**e**). Renilla luciferase activity (Ren-Luc) was used to normalize 8xGTIIC activity. Graphs represent mean ± SEM of 3 biological replicates. **f** Cell viability over indicated time for each cell population as measured by MTT assay (mean ± SD). Shown is representative of three biological replicates with three technical repeats each. **g** RT-qPCR (mean ± SEM) and (**h**) immunoblot analyses of indicated mRNAs and corresponding proteins in U251 cells expressing indicated shRNAs. Student’s *t* test, \**^, #^P* < 0.05, \*\**P* < 0.01, \*\*\**P* < 0.001.

Previously, we generated a chimera construct that consists of a constitutively active Dia1 fragment (*Δ*N3; lacking the auto-inhibitory Rho-binding domain) fused to a C-terminal Syx fragment containing the PDZ binding motif [syx(C)] responsible for membrane recruitment of Syx^8^. Expression of YFP-Dia1(*Δ*N3)-syx(C), hereafter referred to as “chimera”, promotes Dia1-induced signaling events selectively at Syx-targeted membrane complexes and rescues polarity defects in Syx depleted cells^8^. In agreement, expression of this chimera in U251 cells increased YAP/TAZ activity by 5-fold, while expression of YFP-Dia1(*Δ*N3) failed to significantly increase YAP/TAZ activity (Fig. 6e). These data suggest that localized activation of Syx-RhoA-Dia1 signaling promotes the nuclear localization and transcriptional activity of YAP/TAZ to affect gene expression. Confirming this hypothesis, *YAP1* silencing decreased cell growth *in vitro* and altered the expression of the same cell cycle regulators affected by knockdown of *Syx* and *Dia1* (Fig. 6f-h and Fig. S5d).

While the above data indicate a key role for YAP/TAZ in the Syx-RhoA-Dia1 signaling axis, they do not exclude potential contributions from other RhoA-induced signaling pathways. Indeed, depletion of Syx in U251 cells reduced SRF/MRTF activity, both in control cells and under conditions of serum stimulation, as measured by an SRF/MRTF-responsive luciferase reporter (SRF-RE) (Fig. S6a). The growth of GBM cells *in vitro* was also significantly suppressed by CCG-203971, a specific inhibitor of SRF/MRTF signaling (Fig. S6b).

### Targeting Syx-Dia1 signaling cooperates with temozolomide and radiation treatment

The chemotherapeutic agent TMZ improves patient overall survival and is part of the standard of care for GBM along with surgery and radiation therapy^36^. TMZ treatment generates DNA lesions, leading to replication fork collapse, G2/M cell cycle arrest and cell apoptosis, and tumor cells with low or no expression of MGMT are particularly sensitive to this agent. As silencing of either *Syx* or *Dia1* expression increased DNA damage in GBM cells, we postulated that targeting Syx-Dia1 signaling might cooperate with TMZ in suppressing cell growth. To test this hypothesis, we transduced U251 cells with increasing amounts of *Syx*-shRNA carrying lentiviruses, in order to achieve increasing levels of *Syx* depletion (Fig. 7a). After infection, cells were treated with different concentrations (10-300 μM) of TMZ, and cell viability was then assessed using the Cyquant assay (Fig. 7b).

**Fig. 7.**
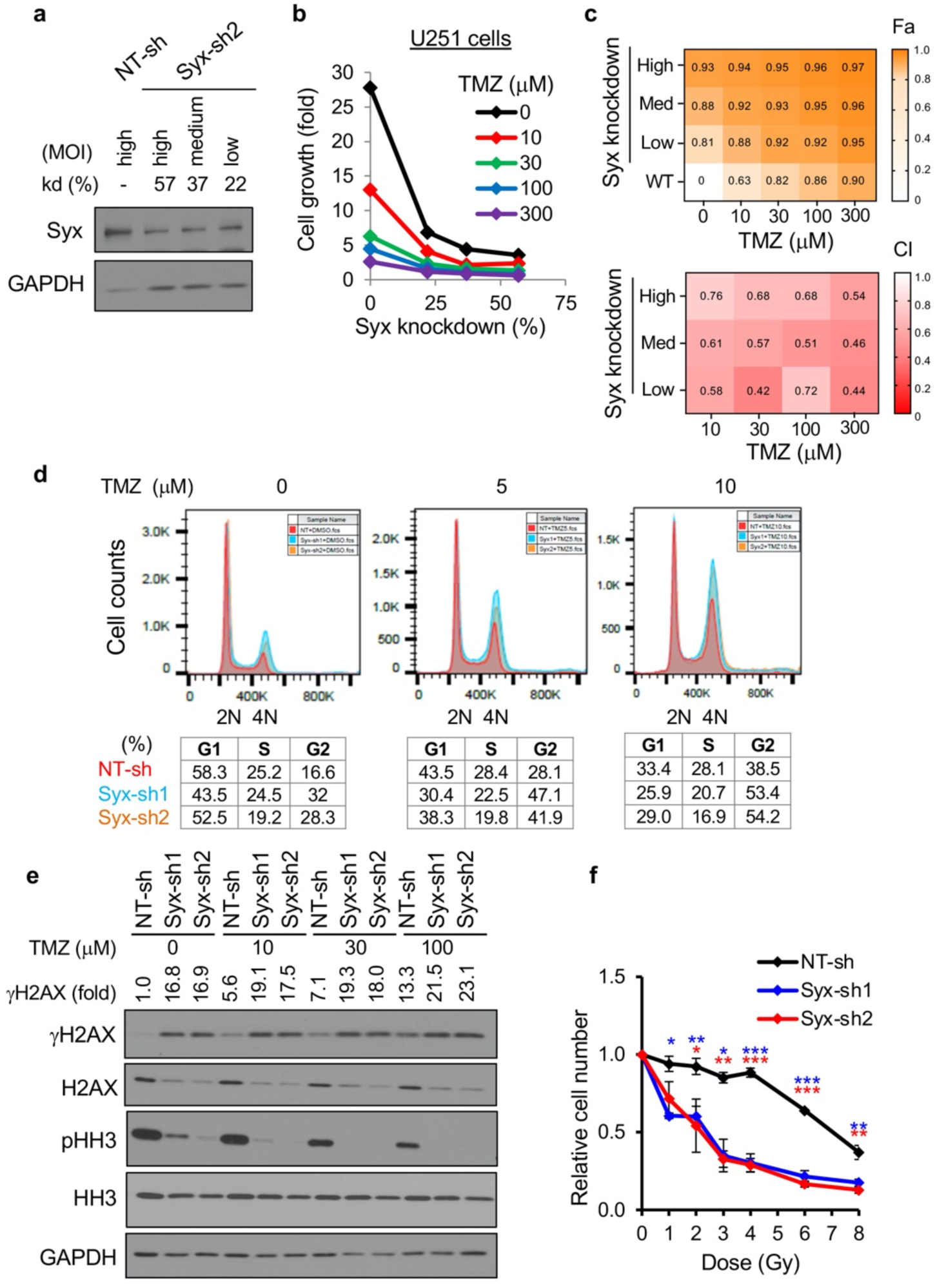
Syx knockdown cooperates with temozolomide and radiation to suppress cell growth. **a** Immunoblot analysis of Syx in U251 cells transduced with Syx-sh2 expressing lentiviruses at different multiplicities of infection (MOI). Syx knockdown efficacy (kd %) relative to GAPDH expression is indicated. **b** Representative graph of 3 biological repeats depicts the growth of U251 cells exhibiting different degrees (%) of Syx knockdown (x-axis) and treated with or without TMZ for 6 days. **c** Heatmaps depict average relative growth inhibition (top) and synergistic interaction (bottom) between Syx targeting and TMZ from 3 biological repeats. Fa, affected fraction. CI, combination index. Orange (top panel) indicates high Fa, which corresponds to high growth inhibition (top). Red (bottom panel) indicates low CI, corresponding to high synergy. **d** DNA fluorescence histograms (top) and cell cycle distribution (bottom) of U251 cells expressing NT-sh (red), Syx-sh1 (cyan), or Syx-sh2 (orange) and treated with 0 μM, 5 μM, or 10μM TMZ. **e** Immunoblot analysis of *γ*H2AX, total H2AX, pHH3, total HH3 and GAPDH in lysates of U251 cells expressing indicated shRNAs and treated with TMZ for 4 days. *γ*H2AX levels normalized to GAPDH and compared to the NT-sh non-treated control are indicated as *γ*H2AX fold. **f** Graph (mean ± SD, 3 biological repeats) shows the relative number of U251 cells expressing indicated shRNAs following treatment with different doses of radiation (Gy). Cell viability was measured 8 days post radiation and normalized to non-irradiated cells. Student’s *t* test, \**P* < 0.05, \*\**P* < 0.01, \*\*\**P* < 0.001.

As expected, knockdown of Syx reduced cell growth (Fig. 7b, black line and Fig. 7c (top panel), column at TMZ=0 μM). TMZ also resulted in a dose-dependent decrease of cell growth in U251 cells, which are sensitive to TMZ due to MGMT promoter methylation (Fig. 7b, datapoints at 0 percent of Syx knockdown and Fig. 7c (top panel), row at WT). Importantly, the combination of Syx depletion with TMZ resulted in a profound and dose-dependent loss of cell viability (Fig. 7b, c (top panel)). We then tested TMZ synergy with Syx targeting and observed a Combination Index (CI) of <1, suggestive of synergistic interaction between the two across all 3 x 4 dose matrix tested (Fig. 7c (bottom panel)). Since both TMZ and Syx depletion cause accumulation of cells in G2/M and DNA damage^3^ (Figs. 2a, 3c-d, and S3a-d), we further assessed the effect of this combination strategy on cell cycle progression, mitotic entry, and DNA DSBs. The combined treatment increased both the extent of G2/M arrest (at low concentrations of TMZ) and levels of DSBs (detected by *γ*H2AX), and decreased mitotic entry (detected by pHH3) (Fig. 7d, e). However, TMZ treatment did not further decrease cyclin levels or increase the level of cleaved caspase 3 induced by *Syx* knockdown (Fig. S7a), suggesting that they act by non-overlapping mechanisms.

As depletion of Syx inhibited the growth of both TMZ-sensitive (GBM12) and resistant (GBM10, GBM14) lines (Fig. 1c and Fig. S1), we postulated that targeting Syx could synergize with TMZ even in lines with acquired TMZ resistance. To test this hypothesis, we utilized a previously established U251 subclonal line selected for TMZ resistance (U251TMZ)^37^. Indeed, we verified that the half maximal inhibitory concentration (IC50) of TMZ for U251TMZ is approximately 10-fold higher than its parental U251 line (58.2 ± 5.1 μM vs. 4.3 ± 1.4 μM, respectively; mean ± SEM) when assayed by the Cyquant method after TMZ exposure. Similar to parental U251 cells, the combination of Syx depletion with TMZ resulted in a profound, dose-dependent, and synergistic loss of viability in U251TMZ cells (Fig. S7b, c). However, TMZ treatment did not further potentiate the effect of Syx knockdown in inducing DSBs or suppressing mitosis entry (Fig. S7d). Overall, these data indicate that targeting the Syx-Dia1 signaling pathway inhibits GBM growth and cooperates with TMZ to induce GBM cell death.

Ionizing radiation is the other arm of the standard of care for GBM. Radiation therapy induces DNA DSBs, leading to cancer cell death. To determine whether targeting Syx signaling can potentiate cell response to radiation treatment, we tested the combined effect of Syx depletion and radiation therapy in U251 cells. Radiation treatment caused a dose-dependent decrease of U251 cell viability, which was significantly increased by concomitant depletion of endogenous Syx (Fig. 7f). The data argue that targeting Syx sensitizes GBM cells to radiation therapy.

Finally, recent studies reported the induction of the endoplasmic reticulum unfolded protein response (UPR^ER^) by DDR^38^, as well as the inhibition of UPR^ER^ by YAP signaling^39^.

Interestingly, Syx depletion promoted the UPR^ER^, as evidenced by the upregulation of key markers including BiP, IRE1*α*, CHOP and JNK phosphorylation (Fig. S7e). The UPR^ER^ cascade can often lead to autophagy for the maintenance of organelle and cellular homeostasis. We observed an increase in LC3 lipidation (LC3-II), indicating the initiation of autophagy by Syx depletion, but no decrease in the lysosomal marker p62/SQSTM1 (Fig. S7f).

## Discussion

Despite multimodal therapy with surgery followed by radiation and TMZ, GBM is a universally lethal disease. Aggressive growth and dissemination of tumor cells in brain parenchyma are major factors contributing to this dismal outcome. The “go or grow” hypothesis argues that cancer cells either migrate or proliferate and based on this, they exhibit different sensitivities to therapies that selectively target migrating or proliferating cells. Therefore, targeting both glioma cell migration and growth may be essential for optimal management of GBM^40^. We previously reported that the Syx-RhoA signaling axis is active in glioma cells, and that suppressing Syx action blocks both random and directed glioma cell migration by reorganizing the cytoskeleton and disrupting microtubule bundling and capture at leading cell edges^8^. Here, we show that activation of Syx also regulates gene expression to promote cell cycle progression and GBM tumor growth. These effects are promoted by the Syx-mediated activation of RhoA, selective activation of the downstream effector Dia1, and at least in part, by YAP/TAZ signaling. Importantly, targeting this pathway cooperates with both radiation treatment and TMZ, the standard of care for GBM. Combined, these data point to a key role for Syx in both GBM growth and migration, and suggest that this pathway can be exploited for GBM therapy.

Upregulation of RhoA expression and activation results in increased cell migration, invasion, and tumor growth^5^. In agreement, our previous work has demonstrated that Syx is part of the Crumbs polarity complex and promotes directed glioma cell migration via its downstream effectors RhoA and Dia1^8, 41^. Increased expression/activation of YAP/TAZ promotes tumorigenesis and correlates with poor clinical outcome in GBM patients^42–45^. Here, we identified a Syx-RhoA-Dia1 signaling axis that regulates gene expression at least in part via YAP/TAZ nuclear signaling to promote glioma cell growth. The mechanism by which Syx-mediated activation of RhoA and Dia1 leads to nuclear localization and activation of YAP/TAZ signaling is unclear, but it could involve Amot, a known YAP/TAZ and Syx interacting partner^18^ ^9^. Notably, YAP/TAZ independent mechanisms may also contribute to the Syx-mediated promotion of GBM cell growth. Indeed, Syx depletion decreased both basal and serum-induced SRF/MRTF-A activity, and treatment of GBM cells with CCG-203971, a specific inhibitor of SRF/MRTF signaling, decreased GBM cell growth (Fig. S6)^45^. Taken together, the data argue for the development of Syx-specific inhibitors. As Syx knockout mice develop normally, we postulate that Syx-targeted therapeutics would be a clinically relevant means of GBM treatment with tolerable toxicity^41^.

Accumulation of DNA damage by multiple mechanisms (radiation therapy, chemotherapy, genetic mutations) triggers DDR, which surveils DNA integrity and induces cell cycle checkpoints and DNA repair mechanisms. Prolonged mitosis is known to cause structural aberrations of chromosomes and DNA breaks^46–48^. Our data show that glioma cells depleted of endogenous Syx are arrested at G2/M, while a fraction of them undergo a prolonged mitosis (Fig. 2d). We postulate that the fraction of cells undergoing mitosis is responsible for the increase in DNA damage, a hypothesis we did not test directly. Both Survivin and Cdc20 are key proteins that ensure proper chromosome alignment and segregation during mitosis^23, 24^. Thus, it is plausible that the prolonged mitosis and the increase in DNA DSBs observed in Syx depleted cells are caused by downregulation of Survivin and Cdc20.

We postulated that the defects in cell division and the resulting DDR induced by targeting the Syx signaling pathway can be exploited for GBM therapy, as they can ultimately lead to cell apoptosis. In support of this hypothesis, combination of Syx depletion and TMZ synergistically inhibited cell growth in both TMZ-sensitive and TMZ-resistant cell lines (Fig. 7a-c and Fig. S7b, c). The data argue that Syx depletion and TMZ engage independent but cooperative mechanisms to promote DNA damage and to inhibit cell growth (Fig. 7a-e and Fig. S7a-d), and suggest a potential role for Syx in TMZ resistance. As both TMZ and radiation therapy induce DDR, we postulated that depletion of Syx may sensitize cells to radiation treatment, a hypothesis that was validated in U251 cells (Fig. 7f).

One concern is that p53 deregulation, which frequently occurs in GBM^49^, can lead to defects in apoptosis upon induction of the DDR. However, we observed similar growth defects by targeting Syx-Dia-YAP/TAZ signaling in either TP53-mutant (U251, LN229, and GBM12) or TP53-wildtype (GBM10, GBM14) cell lines. These data argue that Syx-directed therapeutics could target GBM cells irrespective of their p53 status. We examined the role of Syx signaling in UPR^ER^ and autophagy, two interconnected pathways that can be regulated by both the DDR and YAP signaling and can lead to apoptosis in the absence of p53, if cellular homeostasis cannot be maintained^50, 51^. Syx depletion promoted the UPR^ER^, induced proapoptotic markers CHOP and JNK, and initiated but failed to complete autophagy in our cells, suggesting a loss of cellular homeostasis (Fig. S7e, f). Nonetheless, a recent study reported increased production of autolysosomes, fusion products of autophagosomes and lysosomes, in GBM cells upon Syx knockout^52^. Therefore, while the mechanistic details are still unclear, a potential role for Syx in regulating UPR^ER^, autophagy, and cellular metabolism exists and requires further examination.

In summary, we provide evidence implicating the Syx-Dia-YAP1 signaling pathway in GBM tumorigenesis and uncover its role in gene expression, cell cycle progression, and cell growth. Combined with its ability to promote directed cell migration, the Syx signaling pathway presents a unique target for GBM therapy. This is further underscored by evidence that targeting Syx in combination with TMZ or radiation treatment, the current standard therapy, strongly suppresses GBM cell growth. The clinical translation of these findings is limited by the lack of specific Syx or Dia1 small molecule inhibitors and the poor blood brain barrier penetration of available YAP targeting agents like verteporfin. However, other exchange factors have been targeted, suggesting that specific inhibitors of the Syx-RhoA interaction can be generated^53, 54^. We postulate that targeting Syx, as the most upstream pathway member, will provide maximal efficacy. However, it is possible that other RhoGEFs can also selectively activate RhoA-Dia1-YAP/TAZ signaling in GBM cells. Understanding the nodes of vulnerability to this pathway in different patient tumors will be essential for the development of optimal targeted therapies for this deadly disease.

## Methods

### Study design

Experiments were designed to determine the role of Syx in GBM cell growth, to identify underlying molecular mechanisms of action, and to assess whether targeting Syx signaling can increase GBM sensitivity to chemo- and radiation therapy. Two independent shRNAs for each target gene (Syx, Dia1 and YAP1) and multiple GBM cell lines (2 conventional and 3 PDX lines) were used in the study and cells were maintained for short time periods (approximate 3-4 weeks) to prevent genetic drifting. Cell growth and qRT-PCR assays were carried out with multiple replicates (indicated in figure legends) with three technical repeats. For time-lapse fluorescence imaging experiments in living cells, limited amount of light was used to excite fluorescently labelled proteins to prevent phototoxicity. The survival study *in vivo* was performed in 5 animals per group, which provided sufficient power to assess statistical significance when all animals were included for the determination of survival difference between different groups.

### Constructs, Reagents, and Antibodies

The Hs-Syx and pHIV-H2BmRFP expression plasmids were obtained from GenScript (Ohu22775C, NM_198681 transcript variant 2 mRNA ORF clone) and Addgene (plasmid 18982), respectively. The pEYFP-Dia1(*Δ*N3) (CA-Dia) and pSinLuc constructs were gifts from S. Narumiya, Kyoto University, Kyoto, Japan^55^ and Yasuhiro Ikeda (Mayo Clinic, Rochester)^56^, respectively. The pEYFP-Dia1(*Δ*N3)-syx(C) construct (chimera) was generated previously^8^. Luciferase plasmids, including 8xGTIIC-luciferase (Addgene, plasmid 34615^35^), SRF-RE luciferase (pGL4.34[luc2P/SRF-RE/Hygro], Promega E1350), and human Renilla luciferase (pGL4.74[hRluc/TK], Promega, E6921) were used in the study. The MISSION shRNAs in the pLKO.1 lentiviral vector with a puromycin resistance gene were from Sigma. Product identification numbers for each shRNA are listed – NT-sh: SHC002, Syx-sh1: TRCN0000130291, Syx-sh2: TRCN0000128190, Dia1-sh1: NM_0052192.2-2523s1c1, Dia1-sh2: NM_005219.2-2557s1c1, YAP1-sh1: NM_006106.2-1232s1c1, YAP1-sh2: NM_006106.2-1373s1c1. Chemicals included TMZ (Sigma, T2577), RO-3306 (Sigma, SML0569), and CCG-203971 (Tocris Bioscience, catalogue no. 5277). Antibodies used for immunoblotting and immunofluorescence included: Syx (Abnova, H00057449-M01), GAPDH (Cell Signaling Technology, 2118), *α*-tubulin (Sigma, T5168), pS10-HH3 (Cell Signaling Technology, 9701), HH3 (Cell Signaling Technology, 4499), Cdc20 (Cell Signaling Technology, 4823), Survivin (Cell Signaling Technology, 2808), PARP (Cell Signaling Technology, 9542), cCas3 (Cell Signaling Technology, 9661), pS139-H2AX (*γ*H2AX, Cell Signaling Technology, 2577), H2AX (Cell Signaling Technology, 7631), p27 (Santa Cruz Biotechnology, sc528), p21 (Cell Signaling Technology, 2946), Cyclin D1 (Abcam, ab134175; or Cell Signaling Technology, 2978), Cyclin E2 (Cell Signaling Technology, 4132), Cyclin A2 (Abcam, ab38), Cyclin B1 (Cell Signaling Technology, 4138), Dia1 (BD, 610848), YAP1/TAZ (Santa Cruz Biotechnology, sc-101199), YAP1 (Cell Signaling Technology, 14074), TAZ (BD, 560235), BiP/GRP78 (Cell Signaling Technology, 3177), IRE1*α* (Cell Signaling Technology, 3294), CHOP (Cell Signaling Technology, 2895), pT183/Y185-JNK (Cell Signaling Technology, 9251S), JNK (Santa Cruz Biotechnology, sc-474), LC3-I/II (Acris, AM20212PU-N), p62/SQSTM1 (Cell Signaling Technology, 5114S).

### Cell Culture, Transfection and Transduction

GBM conventional cell lines (U251, LN229, U251TMZ) and PDX lines (GBM10, GBM12, GBM14, provided by Dr. Sarkaria, Mayo Clinic Rochester) were maintained in DMEM (Corning, 10-017-CV) supplemented with 10% Fetal Bovine Serum, 2 mM L-Glutamine, and 1% non-essential amino acids. Penicillin and streptomycin were included in the PDX culture media. U251 cells selected for TMZ resistance (U251TMZ) were established previously^37^. Conventional and PDX lines were maintained in culture for less than 15 and 5 passages, respectively. Transfection was performed using Lipofectamine 200 (Invitrogen) according to the manufacturer’s instructions. RNA silencing in this study was described previously using the MISSION shRNA Lentiviral Transduction (Sigma-Aldrich)^8^. Infected cells were selected with puromycin (2.5 ug/ml for U251, LN229, GBM12, GBM14, and 3.3 ug/ml for GBM10) for 48 hours prior to experiments. The amount of Syx shRNA-expressing viruses was varied for experiments involving TMZ. For mitotic live-cell imaging, U251 cells expressing RFP-H2B were transduced with lentiviral shRNAs followed by puromycin selection as above.

### Animals and Orthotopic Injections

Animals were treated and care-monitored in adherence to the protocol approved by the Mayo Clinic Institutional Animal Care and Use Committee (IACUC). To generate GBM xenografts, 4– 5 week-old female athymic nude mice (Harlan) were anesthetized with isoflurane and short-term explant cultures of GBM12 cells transduced with the luciferase-expression (pSinLuc) and NT or Syx shRNAs lentiviruses were implanted into the brain through intracranial injection, as described previously^57^. Tumor cells (3 × 10^5^ in 5 μl per mouse) were implanted 2 mm lateral and 1 mm anterior to bregma, at 3 mm depth. Animals were monitored daily and maintained until reaching a moribund state. Tumor growth was monitored once a week. Briefly, mice were injected intraperitoneally with luciferin (150 mg/kg/0.1 ml), anesthetized with isoflurane, and imaged with the IVIS Spectrum (Caliper Life Sciences) 10–15 min post-injection. Moribund mice were deeply anesthetized by intraperitoneal injection of 90 mg/kg pentobarbital and euthanized by transcardial perfusion of PBS followed by 4% paraformaldehyde. Brains from mice were resected, cut into four coronal sections of equivalent thickness and fixed in 4% paraformaldehyde overnight at 4 °C. Subsequent formalin fixation and paraffin-embedding (FFPE) was performed and 5 μm-thickness tissue sections were used for the RNAscope assay.

### Immunoblotting

Cells were lysed with RIPA buffer (50 mM Tris, pH 7.4, 150 mM NaCl, 1% NP-40, 0.5 % deoxycholic acid, 0.1% SDS) supplemented with protease inhibitors (RPI, cocktail III) and phosphatase inhibitors (Pierce). In some cases, cells were lysed with 2x Laemmli Sample Buffer (LSB) followed by homogenization through a 29-gauge needle. Protein quantification was assessed using the BCA Protein Assay Kit (Pierce) or the RC DC Protein Assay (Bio-Rad).

Immunoblotting was performed according to standard protocols with ECL (GE Healthcare) reagents. For the synchronization experiment, U251 cells were treated with RO-3306 (9 μM) for 20 hours. Cells at different time points after treatment were collected and lysed in 2x LSB.

### RNAScope

The RNAscope 2.5 HD Brown assay (Advanced Cell Diagnostics) was used to detect mRNAs in 5-μm mouse tissue sections. In situ hybridization was performed according to the manufacturer’s protocol with the Hs-Plekhg5 probe (NM_001042663.1, target region 689 – 1750, cat. 415321). Images were captured using an AT2 slide scanner and ImageScope software (Leica Biosystems).

### Cell Cycle Analysis

Cell cycle progression was determined by flow cytometry using propidium iodine to label DNA content. Briefly, cells at sub-confluency were trypsinized, washed twice with PBS, and fixed in ice-cold 70% ethanol at -20°C overnight. Fixed cells were then incubated with 1mg/ml RNAse A in 0.1% sodium citrate at 37°C for 15 min, and DNA was stained with 100 ug/ml propidium iodide at room temperature for 15 min prior to flow cytometry. DNA content was detected with the Accuri C6 (BD) or the Life Attune NxT cytometers (ThermoFisher). FlowJo and FCS express 5 were used for data analysis. For the combined Syx targeting and TMZ experiment, shRNA lentivirus-infected cells were treated with different concentrations of TMZ for 3 days.

### Cell Growth

Cell growth was assessed using the MTT (Sigma) or Cyquant (ThermoFisher) assays following manufacturer’s protocols. The Cyquant assay was used for experiments involving TMZ. For the MTT assay, cells were plated (U251: 5000 cells, GBM10: 5000 cells, GBM12: 5000 cells, GBM14: 3000 cells per well) in triplicate in 96-well plates and allowed to grow for 5-8 days. For the Cyquant assay, U251 cells were plated (500 cells per well) and allowed to grow for 24 hr prior to 5-6 day treatment with indicated concentrations of TMZ. For the radiation experiments, U251 cells expressing shRNAs were plated (4000 cells per well), subjected to radiation a day later using a RAD-160 Biological Irradiator (Precision X-Ray), and cell viability was assessed by the MTT assay 8 days post radiation. Synergy analysis was performed using the Calcusyn software (Biosoft) with the Chou-Talalay method. Combination index (CI) was used to describe synergistic (CI < 1), antagonistic (CI > 1), or additive (CI = 1) drug interactions. Heatmaps were generated using the GraphPad PRISM software.

### Quantitative RT-PCR

Total RNA was isolated using Trizol (Invitrogen) followed by PureLink RNA minikit (Ambion) according to manufacturer’s protocol. The high capacity cDNA reverse transcriptase kit (Applied Biosystems) was used to convert RNA to cDNA. qPCR reactions were carried out using the TaqMan FAST Universal PCR master mix (Applied Biosystem), in a ViiA 7 or 7900 HT Real-Time PCR system (Applied Biosystems). Data analysis was performed using the RQ Manager (Applied Biosystem) and data is normalized to GAPDH or b-actin. The assay IDs for TaqMan Gene Expression Assay are: GAPDH (Hs99999905_m1), ACTB1 (Hs99999903_m1), Plekhg5 (Syx, Hs00299154_m1), DIAPH1 (Dia1, Hs00946556_m1), CTGF (Hs00170014_m1), BIRC5 (Survivin, Hs00153353_m1), CDKN1A (p21, Hs00355782_m1), CDKN1B (p27, Hs00153277_m1), Cdc20 (Hs00426680_mH), CCNA2 (Hs00996788_m1), CCNB1 (Hs01030103_m1), CCNE2 (Hs00180319_m1), YAP1 (Hs00902712_g1).

### Immunofluorescence and live-cell Imaging

Cells grown on coverslips were fixed in 4% paraformaldehyde/0.12M sucrose/PBS solution at room temperature for 15 minutes, permeabilized with 0.2% Triton X-100/PBS for 5 minutes, and blocked with Protein-Block reagent (Dako, X090930-2) at room temperature for 30 minutes. Proteins were then stained with primary antibodies overnight at 4 °C and then Alexa fluorophore-conjugated secondary antibodies (Invitrogen) for 1 hr at room temperature.

Antibodies were diluted in antibody diluent (Dako, S302281-2). Nuclei were visualized by DAPI (Sigma) staining. Cells were mounted with Aqua Poly/Mount (Polysciences) and imaged using a Zeiss LSM510 META or LSM880 laser confocal microscope under a 40x objective. z-series of images were acquired and maximum intensity projection images were generated before data analysis and are shown in figures. Gamma H2AX (*γ*H2AX) foci and YAP/TAZ subcellular localization were manually analyzed using Image J. For live-cell imaging, images were captured using an Olympus IX83 imaging system equipped with a Stage Top Incubator (Tokai Hit). To examine cell apoptosis in living cells, U251 cells were treated with 4 uM CellEvent Caspase-3/7 green detection reagent (Molecular probes) prior to imaging. U251 cells expressing RFP-H2B and NT- or Syx-shRNAs were plated the day before imaging. Images were acquired every 10 or 20 min over a 24-hour period with a 20x phase objective. Phase-contrast images and RFP-H2B signals were used to identify cells undergoing mitosis. Initiation of mitosis was identified when cells started to shrink and round up and completion of mitosis when two daughter cells were separated. The H2B videos were generated using ImageJ.

### Luciferase reporter assay

U251 cells were plated and transfected the next day with the 8xGTIIC-luciferase and pGL4.74[hRluc/TK] plasmids. The pGL4.74[hRluc/TK] Renilla luciferase construct was used to normalize for transfection efficiency. Medium was replaced 4 hours after transfection. Cells were lysed 24 hours after transfection and the lysates were subjected to dual-luciferase reporter (DLR) assay (Promega) according to the manufacturer’s protocol. Luciferase was detected with a Veritas^TM^ Microplate Luminometer (Turner Biosystems). To assess MRTF-SRF transcriptional activity, U251 cells were transfected with the SRF-RE luciferase and the human Renilla luciferase plasmids. For serum starvation, cells were replated the next day and incubated in serum-free media for 18 hours. Serum stimulation was performed by adding 20% serum to serum-starved cells and incubating for 6 h at 37 °C.

### Statistical analysis

Statistical analysis was performed using the GraphPad Prism software. For in vitro experiments, the unpaired two-tailed Student’s *t*-test was used to determine statistical differences between two experimental groups. For shRNA experiments, statistical comparison between each shRNA of Syx, Dia or YAP1 and NT-sh control was performed. Bar graphs present the data (mean ± SD or mean ± SEM) from multiple experiments or data points. For the animal survival experiment, the Kaplan-Meier method with the Log-rank test was used to compare the survival difference between each Syx shRNA group and the NT-shRNA control. Experimental details are indicated in the figure legends. A *P* value < 0.05 was considered statistically significant.

### Data availability

All data are available in the main text or the supplementary materials.

## Acknowledgments

We are grateful to Ann Tuma (Mayo Clinic, Rochester) for suggestions on the use of TMZ. We also thank Dr. Gasper Kitange (Mayo Clinic, Rochester) for providing the TMZ-resistant U251 cell line, Dr. Arie Horowitz for helpful discussions, and Dr. Steven Rosenfeld (Mayo Clinic, Florida) for his insightful suggestions and critical reading of the manuscript.

## Author contributions

Conception and design: WHL, PZA; development of methodology: WHL, LJL, JNS; data acquisition: WHL, RWF, LMC, LJL; analysis and interpretation of data: WHL, PZA; manuscript preparation: WHL, PZA.

