## Supplemental Information for "A Syx-RhoA-Dia1 signaling axis regulates cell cycle progression, DNA damage, and therapy resistance in glioblastoma"

**One Sentence Summary:** Syx promotes growth and therapy resistance in glioblastoma.

**Authors:** Wan-Hsin Lin[1], Ryan W. Feathers[1], Lisa M. Cooper[1], Laura J. Lewis-Tuffin[1], Jann N. Sarkaria[2], and Panos Z. Anastasiadis[1]*

**Affiliations:** [1]Department of Cancer Biology, Mayo Clinic, Jacksonville, FL32224, USA. [2]Department of Radiation Oncology, Mayo Clinic, Rochester, MN55905, USA. *Corresponding author.

**Supplementary Information**

**Movie S1.** U251 cells transduced with RFP-H2B and NT-sh viruses. Images were acquired every 10 min for 24 hr. Play rate, 10 frames per second.

**Movie S2.** U251 cells transduced with RFP-H2B and Syx-sh1 viruses. Images were acquired every 10 min for 24 hr. Play rate, 10 frames per second.

**Movie S3**. U251 cells transduced with RFP-H2B and Syx-sh2 viruses. Images were acquired every 10 min for 24 hr. Play rate, 10 frames per second.

**Fig. S1**

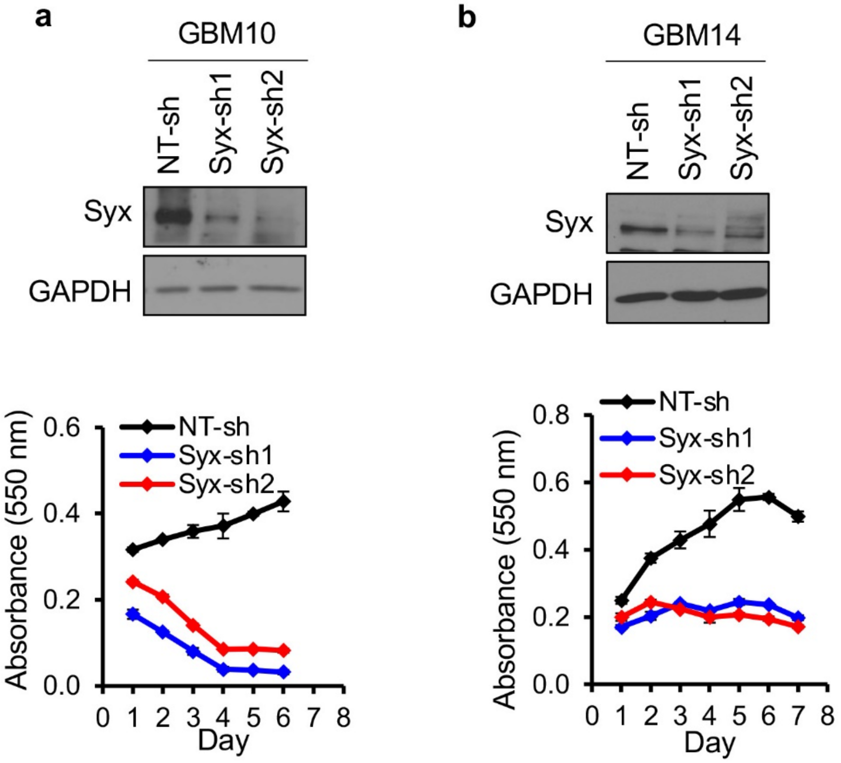


**Fig. S1. Syx knockdown decreases cell growth in GBM PDX lines. a, b** Immunoblot analysis of Syx and GAPDH in lysates from GBM10 (**a**) and GBM14 (**b**) cells transduced with indicated shRNAs (top). Cell viability over indicated time for each cell population was measured by the MTT cell proliferation assay (bottom). Shown are representative graphs with three technical replicates of 3-4 biological replicates. Graphs represent the mean ± SD.

Fig. S2

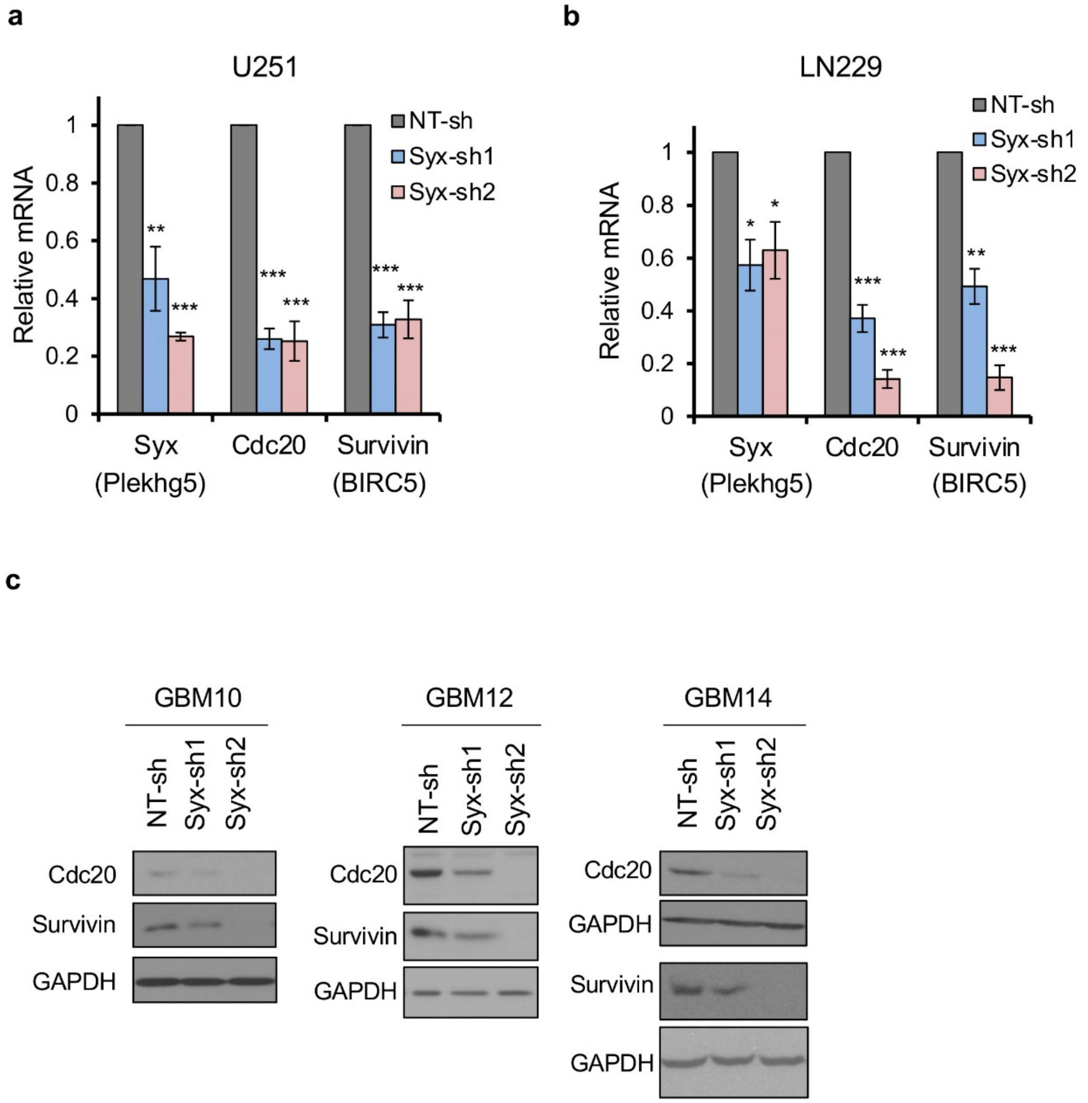


**Fig. S2. Syx depletion downregulates expression of mitosis regulators. a-c** RT-qPCR (**a**, **b**) and immunoblot (**c**) analysis of the expression of mitosis regulators Cdc20 and Survivin (BIRC5) as well as Syx (Plekhg5) in U251 (**a**), LN229 (**b**) cells and three PDX lines (**c**, GBM10: left, GBM12: middle, GBM14: right) that are grown at sub-confluency and expressing indicated shRNAs. Graphs represent the mean ± SEM of at least 3 biological replicates of relative mRNA expression of indicated genes normalized by GAPDH. Student's *t* test was used to assess statistical significance as indicated on graphs. **P* < 0.05, ***P* < 0.01, ****P* < 0.001.

**Fig. S3**

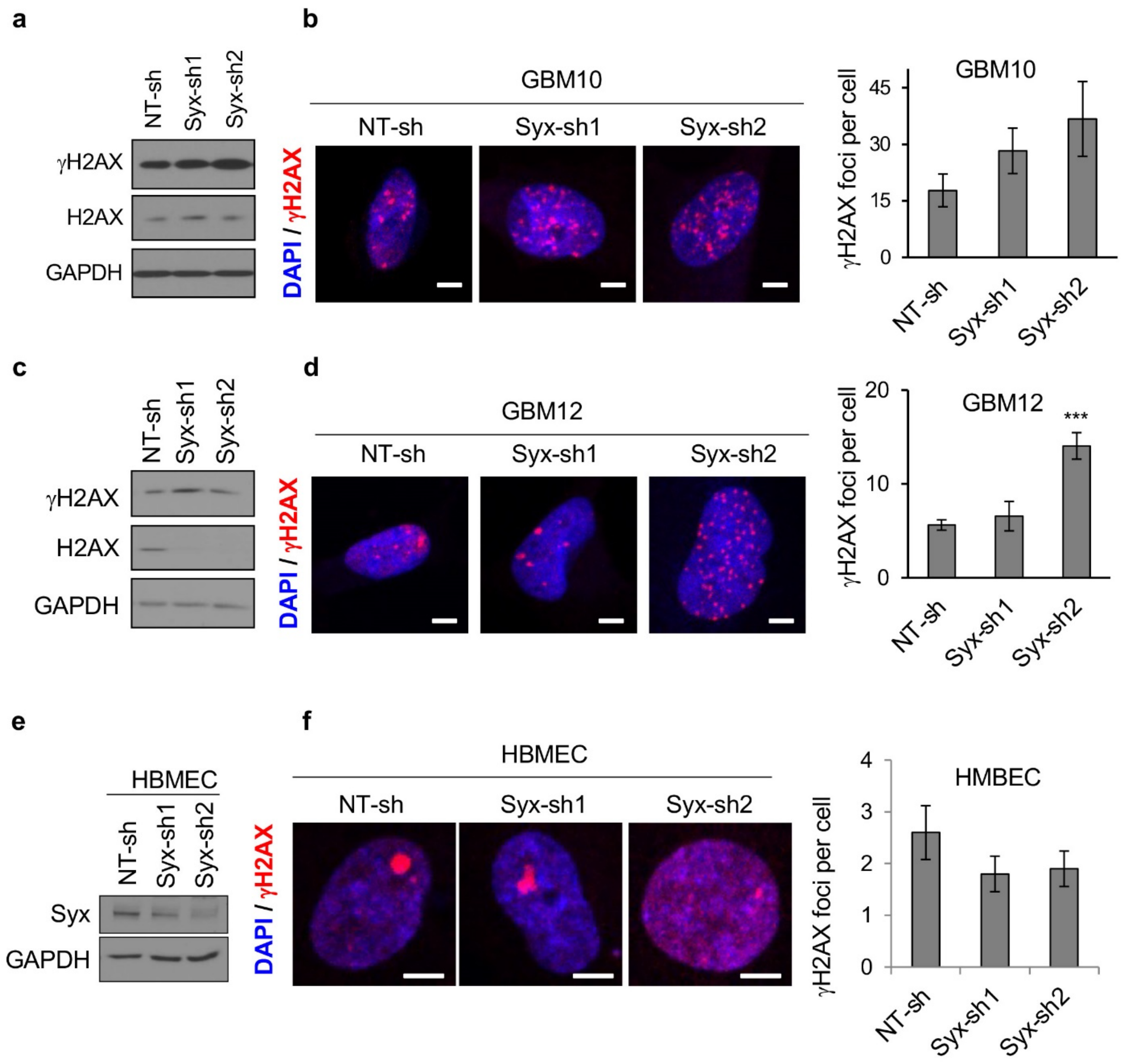


**Fig. S3. Syx knockdown induces DNA damage in GBM PDX lines but not in brain endothelial cells.** **a**-**f** Immunoblot (**a**, **c**, **e**) and immunofluorescence staining (**b**, **d**, **f**) of phosphorylated H2AX at Ser-139 (γH2AX) in GBM10 (**a, b**), GBM12 (**c, d**) and human brain endothelial (HBMEC, **e, f**) cells. Images (**b**, **d**, **f**) show staining for γH2AX (red) and nucleus (DAPI). Scale bar, 5 µm. Shown are maximum intensity projection images. Graphs (**b**, **d**, **f**) on the right represent the average ± SEM number of γH2AX foci per cell. GBM10, *n* = 22-26; GBM12, *n* = 61-62; HBMEC, *n* = 38-45. Student's t-test, $***P < 0.001$.

**Fig. S4**

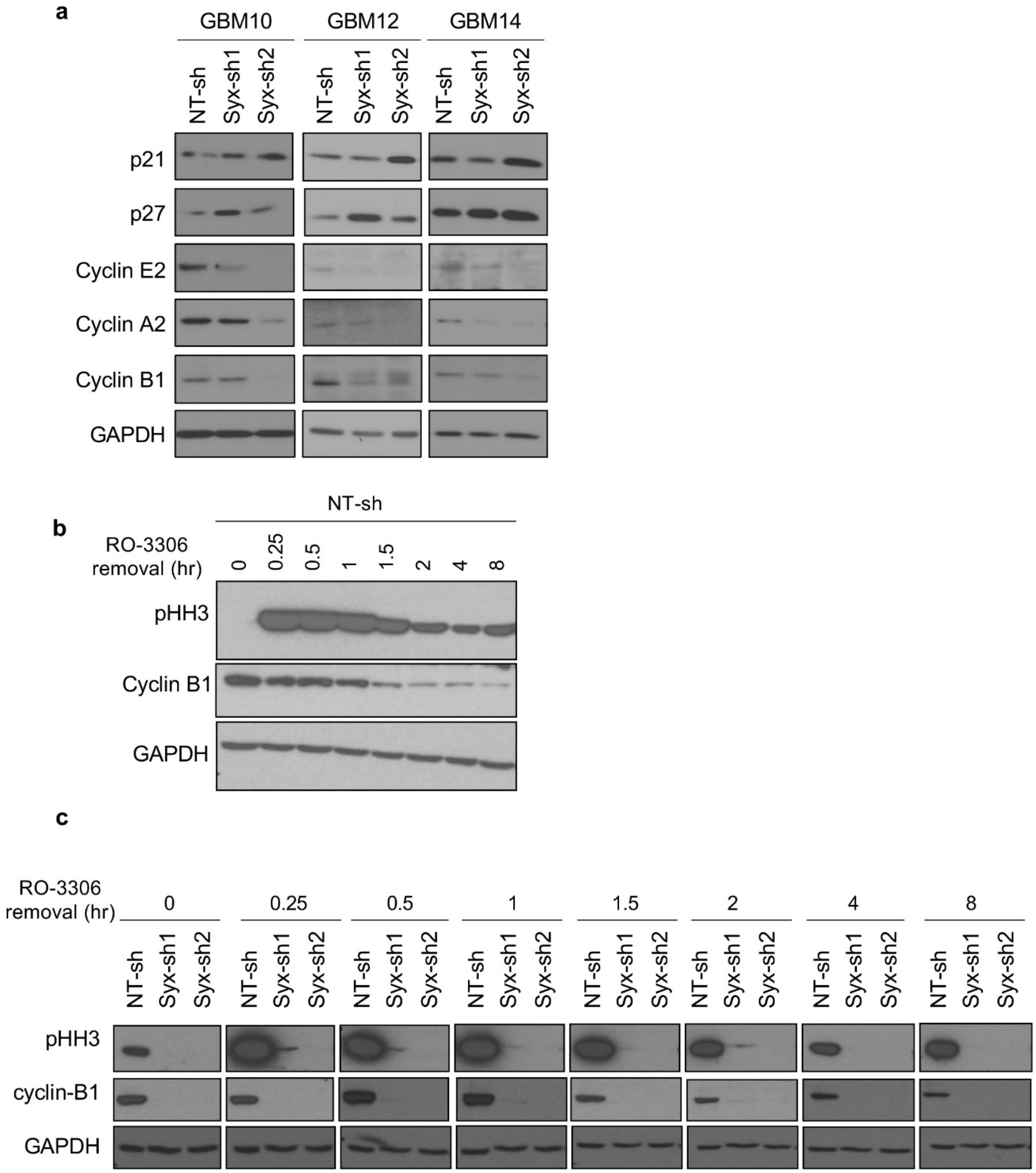


**Fig. S4. Syx knockdown alters expression of cell cycle regulators. a** Immunoblot analysis of p21, p27, Cyclin E2, Cyclin A2, Cyclin B1 and GAPDH in GBM PDX lines. GBM10, left; GBM12, middle; GBM14, right. **b, c** Immunoblot analysis of the levels of phosphorylated histone H3 at Ser10 (pHH3), Cyclin B1 and GAPDH in U251 cells before (0 hr) and after the removal of the G2 phase block RO-3306 (9 μM) at indicated time points. The same lysates from NT-sh (**b**) are also loaded side by side to the Syx knockdown lysates (**c**) for comparison.

Fig. S5

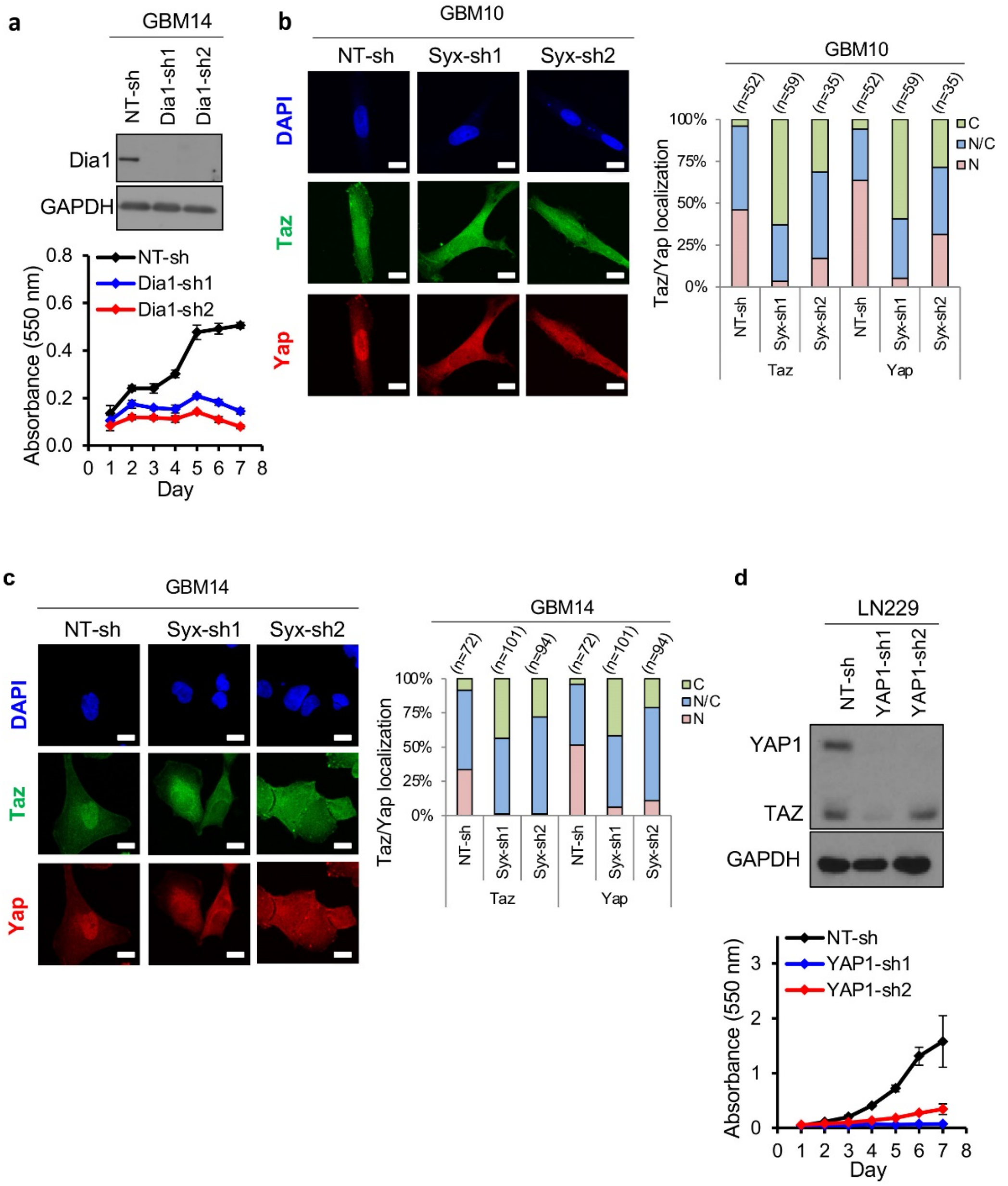


**Fig. S5. Downregulation of the Syx-Dia1 signaling axis reduces cell growth and increases cytoplasmic retention of YAP/TAZ. a** Immunoblot analysis of Dia1 and GAPDH in lysates from GBM14 cells transduced with indicated shRNAs (top). Graph shown is representative of 2 biological repeats and presents cell viability over indicated time for each cell population measured by the MTT assay (bottom). **b**, **c** Immunofluorescence images (left) of subcellular localization of YAP (red) and TAZ (green) in U251 cells expressing indicated shRNAs in GBM10 (**b**) and GBM14 (**c**) cells. Shown are maximum projection images. Scale bar, 15 µm. Staggered graphs (right) depict percentage of cells with YAP and TAZ in the cytosol (C), nucleus (N), or both (N/C). Number of cells analyzed per arm is shown above each bar. **d** Immunoblot analysis of YAP1, TAZ and GAPDH in lysates from LN229 cells transduced with indicated shRNAs (top). Graph shown is representative of 3 biological repeats and presents cell viability over indicated time for each cell population measured by the MTT assay (bottom).

**Fig. S6**

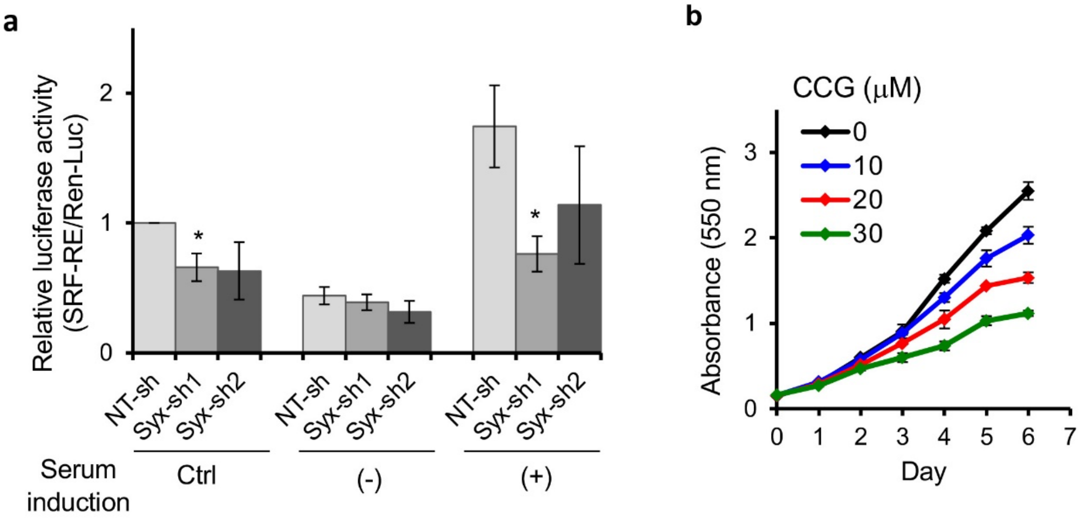


**Fig. S6. Syx knockdown decreases SRF/MRTF signaling. a** Graph shows relative luciferase activity of a SRF responsive promoter containing reporter (SRF-RE) normalized to Renilla luciferase (Ren-Luc) in U251 cells expressing indicated shRNAs. Bar graph depicts luciferase activity of cells grown in regular growth media (10% serum, Ctrl); under serum starvation for 18 h (-), and stimulated with 20% serum for 6 h after serum starvation (+). Bar graphs represent the mean ± SEM of 3 biological repeats with three technical replicates. Student's t-test, *$P < 0.05$. **b** Representative graph of 2 biological repeats performed in triplicate, depicts U251 cell viability following treatment with different concentrations of the MRTF/SRF inhibitor CCG-203971 (CCG) for indicated times.

**Fig. S7**

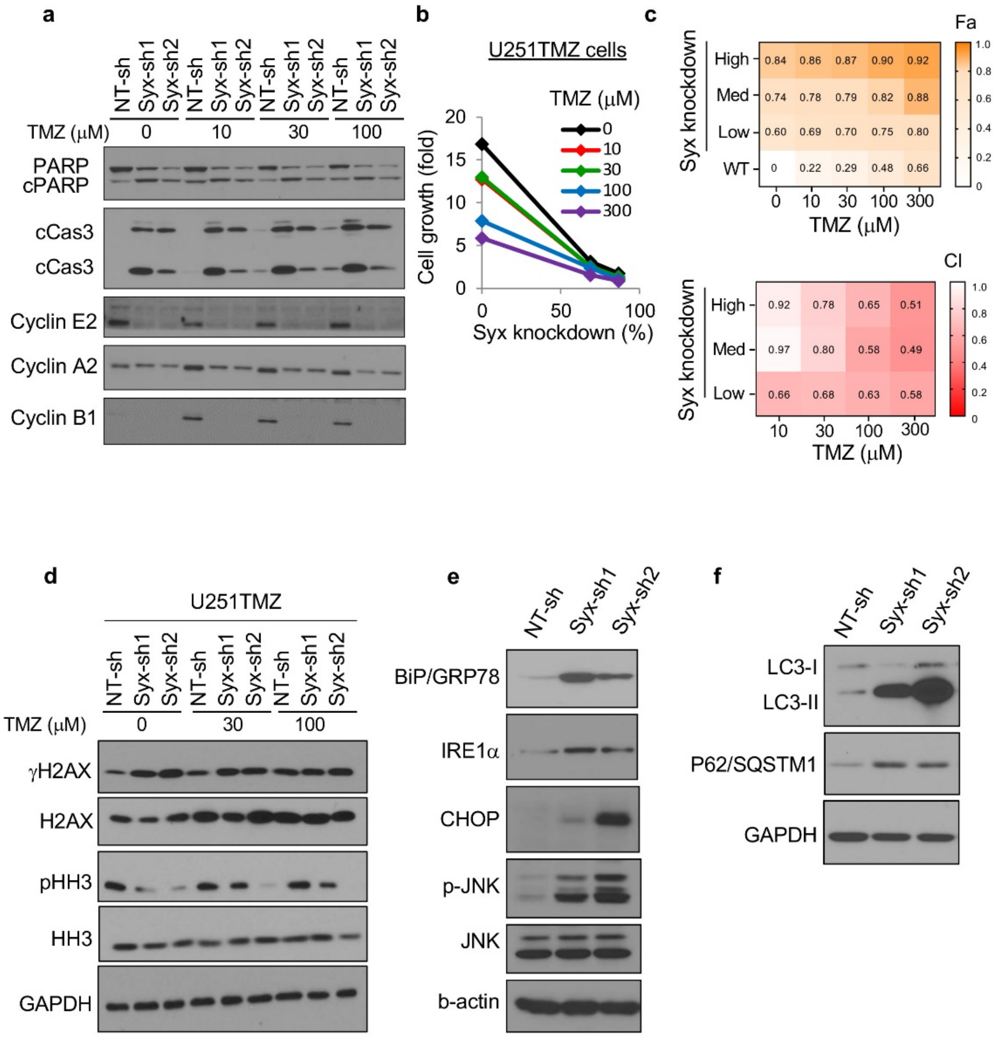


**Fig. S7. Effects of Syx knockdown on TMZ resistance, ER stress and autophagy.** **a** Immunoblot analysis of apoptosis effectors (cPARP, cCas3) and cyclins in lysates from U251 cells expressing indicated shRNAs treated with TMZ at different concentrations for 4 days. **b** Representative graph of 3 biological repeats depicts the growth of U251 TMZ resistant (U251TMZ) cells exhibiting different degrees (%) of Syx knockdown (x-axis) and treated with or without TMZ [0 μM (black), 10 μM (red), 30 μM (green), 100 μM (blue), 300 μM (purple)] for 5 days. **c** Heatmaps present average relative growth inhibition (top) and synergistic interaction (bottom) between Syx targeting and TMZ from 2-3 biological repeats. Different degrees of Syx knockdown were achieved using different amount of Syx-sh2 expressing lentiviruses (Low, 1% virus; Med, 2% virus; High, 4% virus). Fa, affected fraction. CI, combination index. Yellow to white scale (top) indicates high to low affected fraction, which corresponds to high to low growth inhibition (top). Red to white scale (bottom) indicates low to high CI, which corresponds to high to low synergy (CI < 1, synergy). **d** Immunoblot analysis of γH2AX, total H2AX, pHH3, total HH3 and GAPDH in lysates from U251 cells expressing indicated shRNAs treated with TMZ at different concentrations for 4 days. **e, f** Immunoblot analysis of ER stress markers (**e**) and autophagy markers (**f**) in lysates from U251 cells transduced with indicated shRNAs.
